# Osteo-Oto-Hepato-Enteric Syndrome (O2HE) is caused by loss of function mutations in *UNC45A*

**DOI:** 10.1101/208942

**Authors:** Clothilde Esteve, Ludmila Francescatto, Perciliz L. Tan, Aurélie Bourchany, Cécile De Leusse, Evelyne Marinier, Arnaud Blanchard, Patrice Bourgeois, Céline Brochier-Armanet, Ange-Line Bruel, Arnauld Delarue, Yannis Duffourd, Emmanuelle Ecochard-Dugelay, Géraldine Hery, Frédéric Huet, Philippe Gauchez, Emmanuel Gonzales, Catherine Guettier-Bouttier, Mina Komuta, Caroline Lacoste, Raphaelle Maudinas, Karin Mazodier, Yves Rimet, Jean-Baptiste Rivière, Bertrand Roquelaure, Sabine Sigaudy, Xavier Stephenne, Christel Thauvin-Robinet, Julien Thevenon, Jacques Sarles, Nicolas Levy, Catherine Badens, Olivier Goulet, Jean-Pierre Hugot, Nicholas Katsanis, Laurence Faivre, Alexandre Fabre

**Author notes:** These authors contributed equally to this work (Clothilde Esteve, Ludmila Francescatto, Perciliz L. Tan). These authors contributed equally to this work (Aurélie Bourchany, Cécile De Leusse, Evelyne Marinier). These authors contributed equally to this work (Nicholas Katsanis, Laurence Faivre, Alexandre Fabre).

## Abstract

Despite the rapid discovery of genes for rare genetic disorders, we continue to encounter individuals presenting with hitherto unknown syndromic manifestations. Here, we have studied four affected people in three families presenting with cholestasis, congenital diarrhea, impaired hearing and bone fragility, a clinical entity we have termed O2HE (Osteo-Oto-Hepato-enteric) syndrome. Whole exome sequencing of all affected individuals and their parents identified biallelic mutations in Unc-45 Myosin Chaperone A (UNC45A), as a likely driver for this disorder. Subsequent in vitro and in vivo functional studies of the candidate gene indicated a loss of function paradigm, wherein mutations attenuated or abolished protein activity with concomitant defects in gut development and function.

## Introduction

The association of congenital diarrhea and hereditary cholestasis in childhood and infancy is rare, and many patients remain genetically undiagnosed. Recently, some genes identified in one clinical presentation have been expanded to other clinical features. As an example, compound heterozygous and homozygous variants in myosin VB *(MYO5B)* have been identified as a cause of congenital diarrhea with microvillus inclusion disease (MVID; MIM251850) ^1^. Cholestasis, reported as an atypical presentation in MVID, has been considered a side effect of parenteral alimentation. Next generation sequencing approaches have been necessary to span the clinical spectrum to isolated cholestasis^2; 3^. It has been shown that MYO5B, associated with plasma membrane recycling and transcytosis, is essential for polarization of hepatocytes, enterocytes, and respiratory epithelial cells^4; 5^. This broadening of clinical spectra has been important for the demonstration that microvilli are markers of disorders in apical-membrane trafficking and assembly, from bowel to liver ^6; 7^. Such variable clinical presentations have also been demonstrated, for other genes responsible for digestive and liver diseases, including *ABCB11* ^8; 9^, *TTC37* and *SKIV2L* ^10–12^. Here, we investigated four families presenting with a phenotypic constellation that includes cholestasis, congenital diarrhea, impaired hearing and bone fragility. We were intrigued by the fact that, although several of the presenting features are also encountered in various genetic syndromes, we could not identify examples in the literature in which patients shared all observed pathologies. Given these observations, we hypothesized that the constellation of phenotypes in the individuals in our study likely represent a hitherto unknown clinical entity, which we have named tentatively O2HE (Osteo-Oto-Hepato-enteric) syndrome. To understand the genetic basis of this disorder and to provide an entry point towards both mechanisms and potential therapeutic targets, we studied four patients from three families. By combining sequencing with functional studies of candidate genes *in vitro* and *in vivo*, we identified Unc-45 Myosin Chaperone A (*UNC45A*), as a driver for this disorder. UNC45A belongs to the UCS protein family (UNC-45/CRO1/She4p) and has not been linked previously to human genetic disorders. Notably, the *C. elegans* UNC-45 ortholog was first described in a screen for mutations causing motility disorders, a finding likely relevant to the etiopathology seen in humans.

## Material & Methods

### Whole Exome Sequencing (WES)

We obtained written consent from parents under a protocol approved by each of our perspective institutional review boards. Genomic DNA was extracted from peripheral-blood samples from the proband and both parents via standard procedures using the Gentra Puregene blood kit (Qiagen). Family A samples were processed in Dijon University Hospital. The Biobank of the Department of Genetics of La Timone Hospital proceeded with the DNA extraction and ensured the long-term storage of samples from families B and C.

For all patients (A.II.2, B.II.3, B.II.4, C.II.1 and their parents’) whole-exome capture and sequencing were performed at Integragen platform (Integragen SA, Evry, France) from 1.5 µg of genomic DNA per individual using SureSelect Human All Exon V5 kit (Agilent). The resulting libraries were sequenced on a HiSeq 2000 (Illumina) according to the manufacturer’s recommendations for paired-end 75 bp reads. Reads were aligned to the human genome reference sequence (GRCh37/hg19 build of UCSC Genome Browser) with the Burrows-Wheeler Aligner (BWA, v.0.7.6a), and potential duplicate paired-end reads were marked with Picard v. 1.109. The Genome Analysis Toolkit (GATK) v.3.3-0 was used for base quality score recalibration, indel realignment, and variant discovery (both single-nucleotide variants and indels). Read mapping and variant calling were used for data analysis and interpretation. Variants were annotated with SeattleSeq SNP Annotation 138. Variants present at a frequency > 1% in dbSNP 138 and the NHLBI GO Exome Sequencing Project or present from local exomes of unaffected individuals were excluded (see URLs). For the patient A, variants were filtered as described^13^. For patients B.II.3, B.II.4 and C.II.1, variants were filtered with the in-house Variant Analysis and filtration Tool software (VarAFT; varaft.eu). Variant filtering followed the following criteria: i) variants affecting the coding sequence, including splicing and ii) rare variants (<1% allele frequency) from public databases (see URLs), iii) homozygous, compound heterozygous or *de novo* variants. The candidate variants in *UNC45A* were confirmed by Sanger sequencing on a 3500XL Genetic Analyzer^®^ (ThermoFisher, Carlsbad, CA).

### Western blotting

Fibroblasts of patient B.II.3 and all the patients’ lymphoblastoid cells (A.II.2, B.II.3, B.II.4 and C.II.1) were lysed by NP40 cell lysis buffer (ThermoScientific, Rockland, IL, USA) complemented with Halt Proteases and Phosphatases Inhibitor Cocktail 100X(ThermoScientific, Rockland, IL, USA). After incubation on ice for 30 min, lysed cells were sonicated (4 cycles 15s on/15s off), then centrifuged at 16000g for 10min at 4°C. The supernatant was collected and analyzed for protein concentration by Pierce BCA Protein Assay (ThermoScientific, Rockland, IL, USA). Whole lysates of fibroblasts and lymphoid cells (40μg per lane) were loaded onto a SDS-PAGE 10% Bis-Tris gel (Bio-rad, Hercules, CA, USA) for electrophoresis and transferred to a PVDF membrane. The membranes were hybridized with monoclonal mouse antihuman UNC45A antibody (1/1000; ADI-SRA-1800, Enzo Life Sciences, Farmingdale, NY, USA) and with Rabbit anti-human Glyceraldehyde-3-phosphate dehydrogenase polyclonal Antibody (1/10000) (as loading control; Flarebio Biotech llc, USA).

### *In vivo* Zebrafish Assays

All animal work was performed with approved protocols from Duke’s Institutional Animal Care and Use Committee. To determine the effect of *unc45a* suppression in zebrafish, we designed a splice blocking morpholino (MO) against exon 3 of the zebrafish *unc45a* (ENSDART00000159409.1). To assess efficiency of the MO, we extracted total RNA from control and MO injected embryos and performed RT-PCR across the site targeted by the MO, followed by sequencing of the RT-PCR products. For the rescue experiments, we *in vitro* transcribed human WT and V423N-encoding mRNA using the SP6 mMessage mMachine kit (Ambion). All injections were done at the 1-4 cell stage and each experiment was performed in triplicate as described ^14^.

Fluorescent microsphere gavage was performed on 5dpf zebrafish anesthetized in tricaine and mounted in 3% methylcellulose. Injection of Fluoresbrite polychromatic red 2.0 micron microspheres (Polysciences #19508) was gavaged into the zebrafish intestinal lumen as described ^15^. Brightfield and gavage images were taken of 5dpf embryos using the Nikon AZ100 microscope with a 2x objective and a 5.0-megapixel DS-Fi1 digital camera as previously described ^16^.

Serial sections of the zebrafish were obtained by mounting in 3% low-melting point agarose and sections on a vibratome. Standard immunohistochemistry was performed using Phalloidin and 4e8 (both used at 1:1000 dilution). Sections were mounted onto a coverslip using vectashield and imaged on a Nikon Eclipse 90i with a C2-SH camera at 20X. Image analyses and statistics were performed as previously described^17^.

## Results

### Clinical report

All four probands share multiple clinical symptoms including congenital cholestasis, diarrhea, bone fragility with recurrent fractures and deafness (Figure 1; see detailed clinical report in “Supplemental note: Case reports”). They were assembled together after the discovery of independent homozygous or compound heterozygous variants of *UNC45A*, following a genotype-first approach.

**Figure 1:**
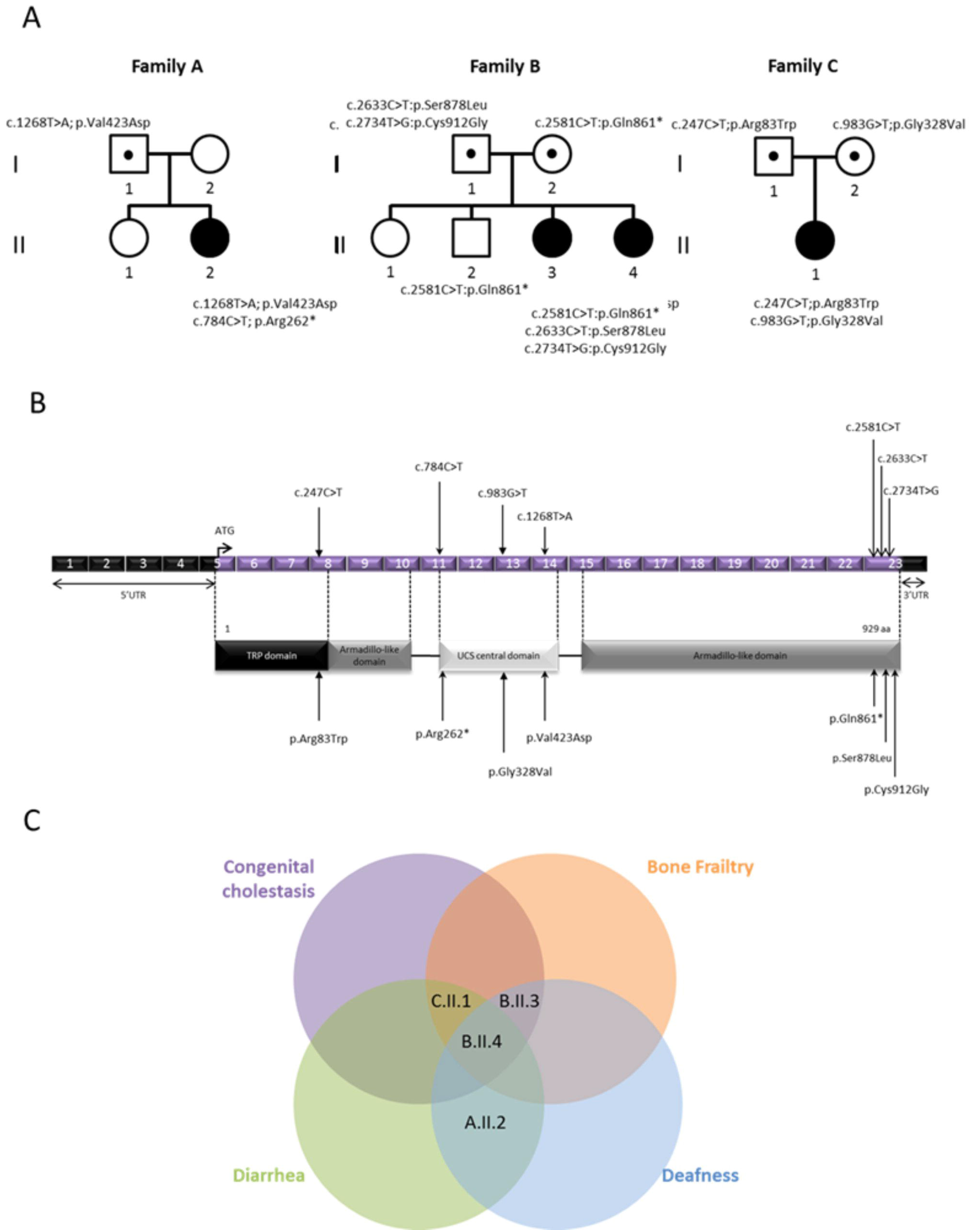
UNC45A variants and gene structure. (A) Pedigrees of the three families with mutations in UNC45A. (B) Structural organization of UNC45A transcript (ENST00000394275) and protein with known conserved protein domains and localization of amino acid residues affected by mutations identified in the three families. Exonic (in purple) regions are not drawn to scale. (C) Venn diagram showing the overlap between clinical signs of the four patients.

#### Family A

The proband is a 5-year-old girl of European descent, the second child of unrelated parents with no family history of digestive disease. She presented intractable diarrhea requiring parenteral nutrition since her fourth day of life. Her evolution was marked by language delay partially linked to severe bilateral perception deafness; she is still with parenteral nutrition. To date, there is no sign of bone fragility or of liver disease.

#### Family B

Two probands are siblings from unrelated parents, originating from Tunisia. There was no known family history of the clinical presentation observed in the two probands. They have one sister and one brother, both of whom are healthy and have normal height. CGH array did not detect any chromosome rearrangements, and targeted sequencing of known disease genes did not reveal any candidate pathogenic variants.

Patient B.II.3, currently 23 years old, presented at 15 days of life icteric cholestasis with normal GGT level. Icterus disappeared at 2.5 years, but elevated levels of bile acid associated with intractable pruritus remained, leading to partial internal biliary diversion at the age of 19 allowing an amelioration of the pruritus. She presented bone fragility: 23 fractures with normal levels of PTH and vitamin D. Severe bilateral perception deafness was diagnosed in her fifth year. She presented a failure to thrive with a current weight of 38.5 kg and height of 147.5 cm and a slight intellectual disability. (See full report in Supplemental Data).

Patient B.II.4, currently 18 years old, presented an icteric cholestasis with elevated GGT at 7 days of life. Like her sister, icterus resolved at 3 years, but cholestasis with elevated level of GGT remained as well as elevated levels of bile acid associated with intractable pruritus. These issues led to partial internal biliary diversion at the age of 12, allowing an amelioration of the pruritus. She had bone frailty with multiple fractures and osteonecrosis of the femoral head at the age of 14, secondary to left hip dysplasia at birth and normal calcium phosphate, D vitamin a PTH levels.

Perception deafness was discovered in her teens. Initially, she presented diarrhea requiring parenteral and enteral nutrition. Diarrhea resolved with time, but a failure to thrive persisted at the last evaluation with a weight of 38.5kg (−3SD) and a height of 147.5cm (−2.5SD) and a mild intellectual disability was noted like her sister.

#### Family C

The proband is a 5-year-old female born to unrelated parents of Turkish origin. The father is carrier of a polyglobulia. After 15 days of life, the proband presented with cholestatic icterus with normal level of GGT associated with liver failure; both resolved at 4 months of life. She presented bone fragility with two spontaneous fractures, with normal calcium phosphate, PTH and D vitamin levels. She had diarrhea since the age of 2 months, requiring parenteral nutrition. She is still with parenteral nutrition and has had recurrent episodes of cytolysis.

### Whole Exome Sequencing of family A and B identified variants in UNC45A

We adopted a genetic approach for the study of a case of intractable diarrhea. The sporadic case in Family A, was found in a large cohort of patients without genetics diagnosis. This study was part of an “undiagnosed program” focusing on the identification of the genetic basis of pathologies in more than 500 affected individuals.

The exome sequencing of this family was first analyzed following a trio strategy. The autosomal *de novo* hypothesis allowed the identification of a *de novo* c.1268T>A; p.Val423Asp variant in *UNC45A*. A second analysis under an autosomal recessive hypothesis identified nine genes with homozygous or compound heterozygous variants in their coding region. *UNC45A* and *CLRN2* were considered the strongest candidates: neither has been associated to date with a clinical phenotype in human disease and only the variant in *UNC45A* c.784C>T; p.Arg262* encodes a nonsense allele. Among the remaining genes, two have been associated with a human pathology: *FRAS1* [MIM: 607830] and *MYO15A*. Frasier syndrome can be likely excluded due to the divergent clinical presentation, while *MYO15A* formally remains a candidate for the deafness phenotype of the patient. However, there are no functional clues about the variant in *MYO5B*, which is thus classified as a variant of unknown significance (VUS). Sanger sequencing for the discovered *UNC45A* mutations confirmed that the missense variant was inherited from the father and the stop-gained mutation was *de novo* (compound heterozygous status).

Independently, we performed whole exome sequencing on genomic DNA from individual B.II.3 and B.II.4 and their parents B.I.1 and B.I.2 (Family B; Figure 1). The trio-based analysis did not identify likely clinically relevant variants in known disease-associated genes. Under the hypothesis of either an autosomal recessive or *de novo* dominant mode of transmission, a search for homozygous, compound heterozygous or *de novo* rare variants (<0.1% in ExAC Browser and GnomAD) allowed us to identify a unique mutated gene shared by the two sisters: *UNC45A*, a locus not yet associated with any known human genetic disorder. Our exome findings and the high predicted expression levels of *UNC45A* in bone, colon and liver (Proteomics DB), rendered this locus a strong candidate. The three identified *UNC45A* variants were confirmed by Sanger sequencing and their segregation was consistent with an autosomal recessive trait (Fig. 1). The two missense c.2633C>T; p.Ser878Leu and c.2734; p.Cys912Gly and the nonsense c.2584C>T; p.Gln861* mutations are all predicted to be damaging by UMD Predictor^18; 19^. They are not found in gnomAD database, except for the variant c.2633C>T; p.Ser878Leu which is reported seven times in the heterozygous state (7/277162).

Biallelic *UNC45A* variants were the only sites shared among the two unrelated families with overlapping phenotypic features, thereby bolstering our interest for a likely causal association of *UNC45A* variants with the disease presentation.

### Additional families with phenotypic match: discovery of family C

We added *UNC45A* to our candidate gene list and we reanalyzed the exomes of families without molecular diagnosis with phenotypes similar to those exhibited by individuals of families A and B. We discovered two missense mutations in a third unrelated family presenting liver disease and bone frailty. Specifically, in family C, we identified two missense mutations c. 247C>T; p.Arg83Trp and c.983G>T; p.Gly328Val in *UNC45A* (Figure 1). These substitutions are predicted to be damaging; one is only found four times (frequency of 4/276948) in heterozygotes, whereas the second one has never been reported in gnomAD Browser. Given the overlap of clinical features of individual C.II.1 with the clinical spectrum of the other patients described here, these results provided further evidence for a causal association of biallelic *UNC45A* mutations with this phenotypic presentation.

### Functional impact of variants found in UNC45A on protein level

To investigate whether these alleles have an effect on protein abundance, we performed Western blot using a monoclonal antibody against UNC45A in all the lymphoblastoid cell lines available in our cohort. We detected UNC45A at the predicted size of 103KDa; quantification of the observed signal in each patient (and the parents of the patient C.II.1) showed a decrease of UNC45A abundance across all patients’ lymphoblastoid cells. Specifically, a significant decrease of protein abundance in each of, A.II.2 B.II.3, B.II.4, C.II.1, (80%, 93%, 89%, 70%, decrease respectively) compared to control cells (Figure 2 C, D). Furthermore, we did not detect any signal at ~70KDa, which would have corresponded to the truncated protein translated from the *UNC45A* allele carrying the nonsense mutation p.Gln861*, suggesting that the truncated protein product is unstable. Regarding nonsense variant p.Arg262* in patient A, which maps to exon 11, we likewise did not detect any truncated protein. However, we cannot formally exclude the possibility of a truncated protein since we do not know the lowest detection threshold of our assay; this data suggests there is nonsense mediated mRNA decay (NMD). Of note, in Family C, father C.I.2 who is heterozygous for p.Arg83Trp and the mother C.I.2 who is heterozygous for p.Val423Asp, also had reduced UNC45A levels (45%, and 52% respectively), also indicating that these alleles are associated with reduced protein expression level and lead to the abrogation of protein stability. Together, all these studies suggest a loss of protein abundance/activity paradigm and thus a loss of function disorder.

**Figure 2:**
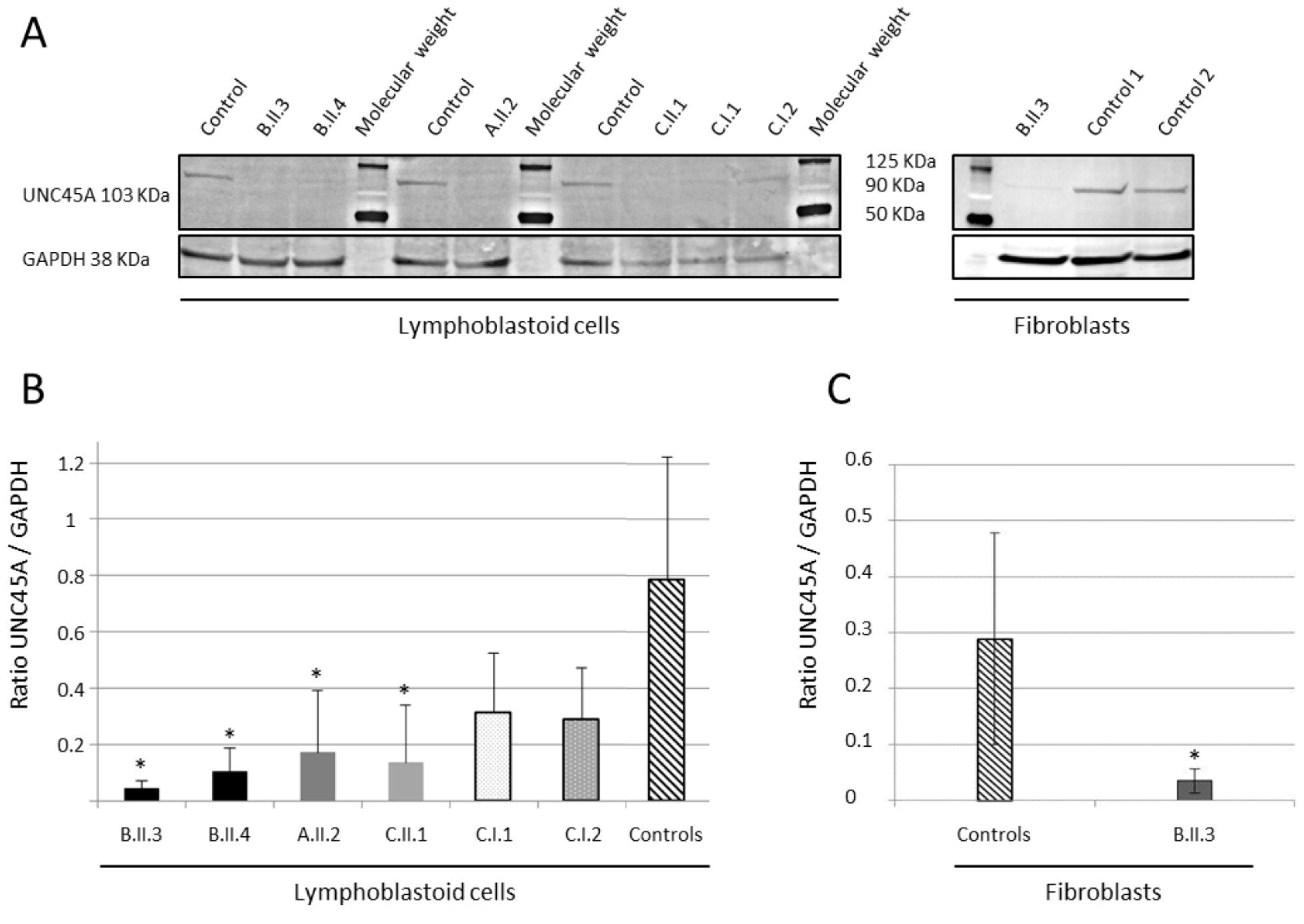
Western blot detection and quantification of UNC45A protein in controls and patient cells. (A) Detection of UNC45A by Western blot analysis in lymphoblastoid cells and fibroblasts using a monoclonal antibody. GAPDH was used as a loading control. Quantification of UNC45A protein levels with Fidji software shows significantly lowered levels of the expression of UNC45A in all patients’ lymphoblastoid cells (B) and fibroblasts of patient A.II.3 (C). Values are mean ratio +/− SD, n=3; Welch’s two sample t-test (patient vs control) (* p-value<0.04).

We next asked whether the reduction of UNC45A expression was restricted to lymphoblastoid cells. Using whole cell lysates from patient B.II.3 fibroblasts, we observed an 82% decrease of UNC45A compared to control cells (Figure 2 A, C). Similar to the previous experiment, we could not detect any signal that might represent the truncated protein, again suggesting NMD. Quantification of the *UNC45A* mRNA produced in these cells was consistent with this notion; we observed a 40% and 50% decrease of transcripts levels in lymphoblastoid cells of patients B.II.3 and B.II.4 respectively (Figure S3).

### In vivo functional testing of the UNC45A variation in zebrafish

With the identification of *UNC45A* likely loss of function mutations in patients with O2HE syndrome, we next sought to ask whether dysfunction of this gene contributes to experimentally tractable aspects of the phenotype. We have shown previously the utility of the zebrafish system to assess the involvement of genetic lesions in the formation and function of the gut ^14; 20; 21^. The zebrafish locus is predicted to encode three splice isoforms of *unc45a*; we focused our studies on the canonical (longest) isoform (ENSDART00000159409.1) which is 64% identical to human UNC45A (ENSG00000140553); the other two isoforms are shorter (230aa and 218 aa versus 935 AA for the canonical isoform) (Figure S8 and S9). We designed a MO against the splice donor site of zebrafish *unc45a* exon 3. This resulted in the inclusion of intron 3, leading to a premature stop.

To assess one of the primary clinical phenotypes of our patients, congenital diarrhea, we tested the effects of suppression of *unc45a* on both enteric neurons and intestinal motility. We first asked whether this phenotype could be caused by a neurodevelopmental enteric defect. We therefore visualized and counted the neurons along the zebrafish gut by staining with antibodies against HuC/D. In triplicate experiments performed blind to injection cocktail, we could not detect any appreciable difference in the number of enteric neurons in morphants versus controls, suggesting that *unc45a* suppression does not impact neurodevelopment in the gut (data not shown). Next, we assayed intestinal motility by fluorescent microsphere gavage into the anterior intestine of 5 dpf (days post fertilization) embryos ^15; 16^, followed by recording of the rate of intestinal motility of the microspheres as a function of time (imaging at 0, 3, 6, 9 and 24 hrs). At each time point we scored in which intestinal zone (1-4, based on anatomical landmarks) the fluorescent microspheres were located as a measurement of transit through the intestine post-gavage. Performing ordinal logistic regression and repeated measures analysis, we saw that, as time progresses, control larvae are less likely to have microspheres remaining in their intestinal lumen compared to morphants (p<0.0001; OR=0.504), with a majority of morphants having microspheres remaining in their intestine after 24 hours post gavage. These data suggest that suppression of *unc45a* leads to an intestinal motility defect (Figure 3).

**Figure 3:**
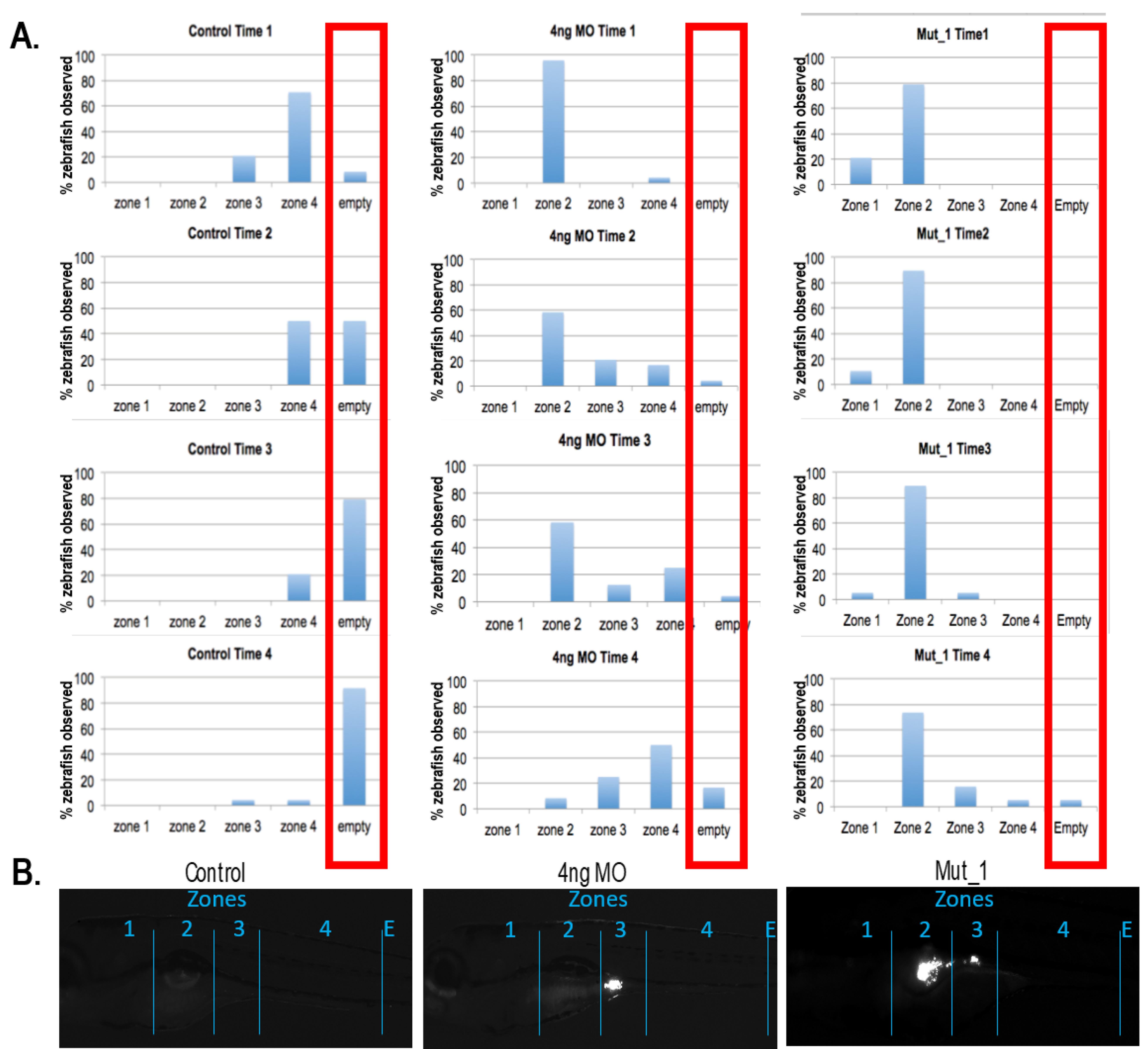
Microgavage assay to assess intestinal motility. Fluorescent microspheres were gavaged and their transit was observed at 0, 3, 6, 9 and 24 hrs. Both morphants and mutant zebrafish for unc45a show intestinal mobility defects.

To test this observation further, we turned our attention to a stable genetic model. We obtained the kurzschlusstr12 (kus^tr12^) zebrafish mutant, an aortic arch mutant identified previously in a forward genetic screen caused by a nonsense mutation in *unc45a* ^22^. Although gut motility phenotypes had not been reported previously, such phenotypes are not readily observable; we therefore tested mutants directly by gavage of fluorescent pellets, followed by measurement of the clearance efficiency of these along the alimentary canal as a function of time. Specifically, we divided the gut into four anatomically-defined zones and we measured the proportion of embryos that had GFP present in each zone. Consistent with our exome findings, kus^tr12^ mutants had a significant delay in clearing fluorescence that was even more pronounced than what we observed in our morphants. Repeated measures analysis showed that as time progresses, control larvae are less likely to have microspheres in their intestinal lumen (the majority of had no microspheres in their intestines) compared to kus^tr12^ fish (p=0.0009; OR=0.1387) with the majority of the microspheres in the kus^tr12^ larvae remaining in zone 2 (compared to morphants that showed accumulations primarily in zones 3 and 4 (Figure 3). The observed motility defect was not due to dysfunctional peristalsis, as there was no appreciable peristalsis defect in kus^tr12^ mutants (data not shown).

We next asked if the intestinal motility defect could be linked to structural defects. Observations of control and mutant animals under brightfield showed a lack of folds in the intestine of kus^tr12^ fish. We therefore performed serial sections and used markers for F-actin (Phalloidin) and intestinal brush borders (4e8) to determine if the defects in intestinal motility we observe in the mutant larvae were the result of structural abnormalities. Analysis of kus^tr12^ mutant and control serial sections, revealed defects in the structure of zone 2 and 3 of kus^tr12^ animals (Figure 4). Staining with Phalloidin revealed that, as opposed to having the expected epithelial folds lining the lumen, kus^tr12^ mutant intestinal tubes are devoid of folds in zone 2 and 3, corresponding to the anterior intestine. The differences in structure were restricted to, and specific for, zones 2 and 3; there were no structural differences in zone 4 and no brush border defects observed in the kus^tr12^ embryos, as their enterocytes maintained the expression and localization of 4e8, the absorptive cell marker.

**Figure 4:**
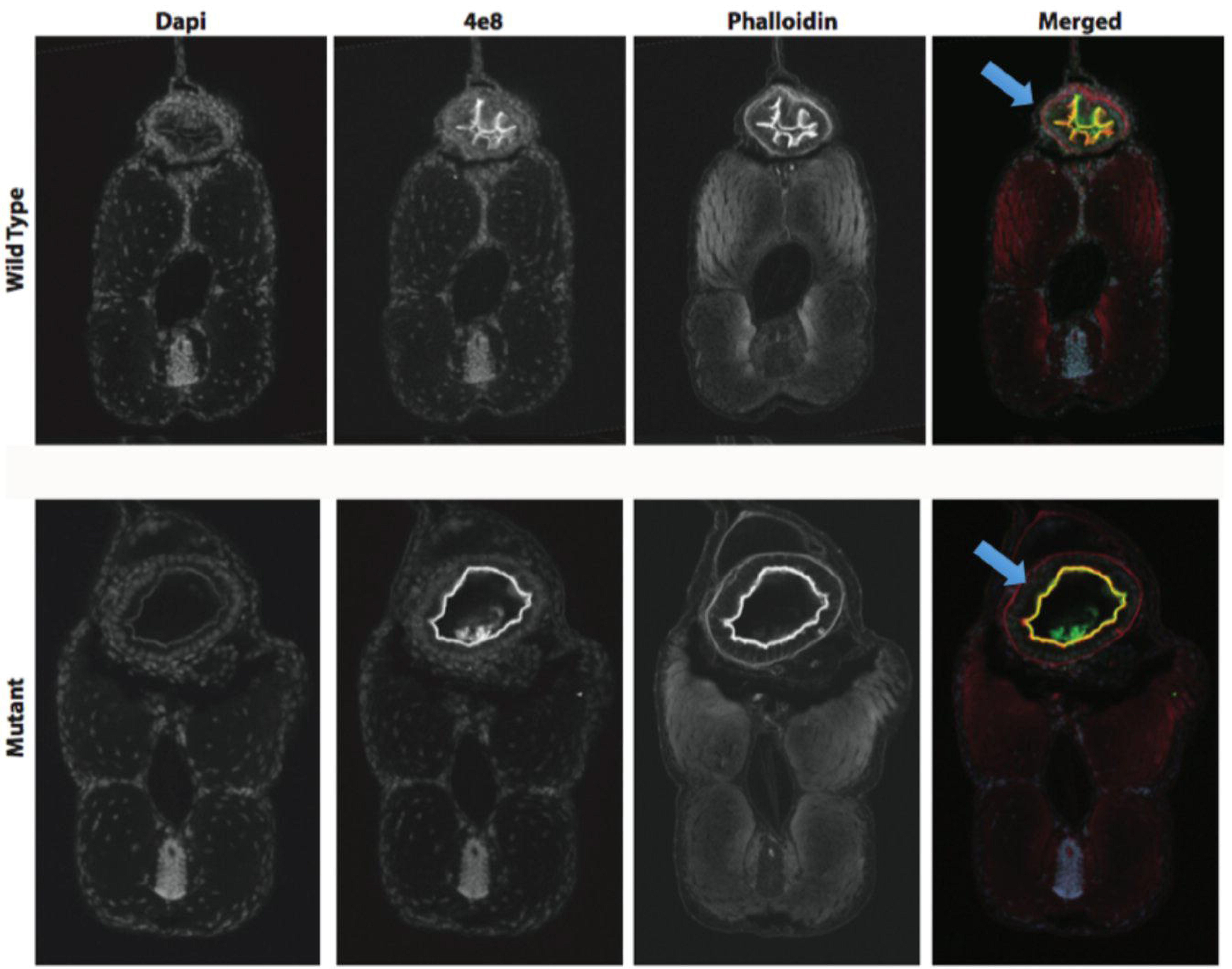
Histological study of transverse sections of zones 2-3 of 5dpf zebrafish. Markers for F-actin (Phalloidin) and intestinal brush borders (4e8) were used and revealed defects (lack of epithelial folds in the intestinal tube) in the structure of kus^tr12^ larvae compared to controls.

Given these phenotypes, we proceeded to test the functionality of the alleles discovered in our cohort. In the Family A, the proband was a compound heterozygote for p.Arg262* and p.Val423Asp, the first mutation being considered to encode a null allele. We therefore modeled p.Val423Asp. To reduce the phenotypic variability that we have sometimes found in morphants, we focused our studies on the of kus^tr12^ homozygous mutants. Specifically, we injected both human wildtype (WT) and V423N encoding mRNA into kus^tr12^ embryos at the 1-4 cell stage and we assessed their ability to improve the structural phenotypes observed in zone 2 of 5dpf mutant guts. Injection of WT human mRNA was able to ameliorate the observed fold defects. In contrast, human p.Val423Asp mRNA injected kus^tr12^ larvae were 3.44 times more likely to have no folds in zone 2 than kus^tr12^ mutants injected with WT mRNA (Figure 5, p=0.005; OR=3.441), with modest overall amelioration of the mutant phenotype. Human WT mRNA was unable to rescue zone 3, likely because it forms somewhat later in development at which point the injected mRNA is mostly extinguished, while zone 4 gave no pathology in mutants and therefore served as an indirect quality control for specificity. Taken together, these data suggested that UNC45A plays a role in the development of a functional intestinal system and that the *UNC45A* variant p.Val423Asp identified in family A likely results in minimal (but not fully extinguished) UNC45A activity.

**Figure 5:**
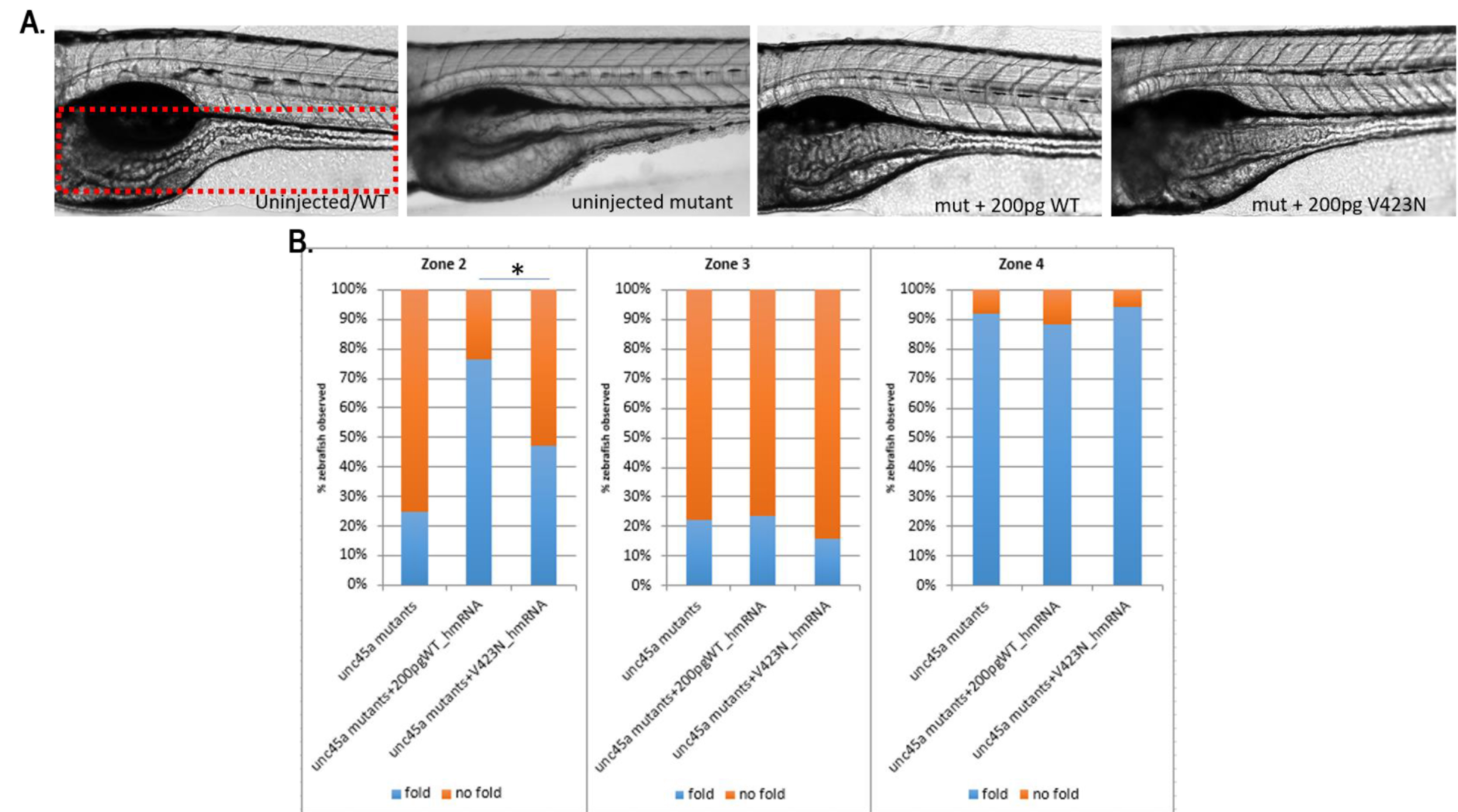
Assessment of human UNC45A c.1268T>A, resulting in a p.Val423Asp change (V423N), on zones 2-3 of mutant unc45a zebrafishes. Mutant embryos were injected with either WT or V423N encoding mRNA and intestinal folds were observed using brightfield microscopy. Mutants injected with WT encoding mRNA were able to restore folds in zones 2-3 however those injected with V423N encoding mRNA were not restored, illustrating that V423N is pathogenic.

## Discussion

We report a hitherto undescribed constellation of phenotypes that we propose to represent a distinct clinical entity, Osteo-Oto-Hepato-Enteric syndrome (O2HE). The main clinical features of O2HE syndrome include congenital diarrhea, congenital cholestasis, bone fragility and deafness. In our families’ exomes, no mutations were found in genes previously known to be involved in cholestasis, brittle bones, or chronic diarrhea. Together, our *in vitro* and *in vivo* data provide strong evidence that loss of UNC45A function results in a previously unpublished clinical pathology, that includes loss of normal digestive intestinal transit.

Based on current knowledge, there is no obvious link between the clinical expression observed in the affected individuals in the three families and UNC45A function. Most studies of *unc45* have been performed in *Caenorhabditis elegans.* However, invertebrates have a single *unc-45* gene that is expressed in both muscle and non-muscle tissues, whereas vertebrates possess one gene expressed in striated muscle (*UNC45B*) and another that is expressed more ubiquitously (*UNC45A*). In fact the phylogeny of UNC-45 homologues showed that this protein family appeared in the Holozoa lineage (Figure S4) and has undergone a duplication event in an ancestor of Vertebrates. This event allowed the specialization of the two resulting paralogues: the muscle Unc45B and ubiquitous Unc45A. This divergence might explain the absence of skeletal muscle pathology in the three families studied here, although additional affected individuals will be required to define the full range of the clinical pathologies associated with *UNC45A* mutations in humans.

UNC45A belongs to the UCS protein family (***U***NC-45/***C***RO1/***S***he4p). *C. elegans unc-45* was first described after a screening for mutations causing motility disorders (UNC stands for uncoordinated) ^23^. The UNC-45 protein has three recognizable domains: an N-terminal tetratricopeptide repeat (TRP) domain (~115 amino acids); a central domain of ~ 400 amino acids and a C-terminal UCS domain (~400 amino acids) ^23; 24^. The TPR domain participates in protein-protein interactions, especially with Hsp70 (HSPA1A [MIM 140550]) and Hsp90 HSP90AA1 [MIM: 140571]) ^25^. The role of the central domain remains unclear, while the C-terminal UCS domain is critical for myosin binding ^24^. Whether or not variants located within specific domains of UNC45A lead to different functional outcomes is still unknown. UNC45A appears to be ubiquitously expressed and has been postulated to be involved in cytoskeletal functions, such as cell division or exocytosis ^26; 27^. These data are consistent with the multi-organ defects observed in our affected individuals, as well as the structural pathology in the gut of our zebrafish mutants. However other features of the protein are not reflected in the phenotype seen in humans. For example, UNC45A co-localizes with type II muscle myosin heavy chain B as well as type V myosins and plays an essential role in myoblast fusion and cell proliferation ^28–31^. However, none of our affected individuals present muscular alterations; none have muscle weakness and all present normal Creatine Phosphokinase level. Recently, *in vivo* studies have reported that *unc45a* plays a role in aortic arch development and could be one underlying cause of human vessel malformations ^22^. This, coupled to reports that individuals with visceral arteriovenous malformations can be more susceptible to cholestasis ^32^. makes UNC45A an attractive candidate; however none of the people in our study display vessel malformation. Although this might be due to differences in development and physiology between lower organisms and humans, we favor the hypothesis in which minimal activity of the human alleles (in contrast to null homozygous zebrafish embryos) might protect from this pathology. The discovery of additional mutations at this locus will help clarify these questions.

There is a core phenotype: four main signs seen at least in three patients (Figure 1, C). Phenotypic variability is observed in the three families studied here; however this is seen in other congenital diseases (for example Microvillus Inclusion Disease or Tricho-Hepato-Enteric Syndrome) and expected in such complex diseases. The four patients present some degree of developmental delay but, we can’t exclude that it is a non-specific symptom. Indeed in the patients A.II.2, B.II.3 and B.II.4, the delay could be a secondary effect of the deafness. There are also some isolated signs, such as the tubulopathy and poor glycemic regulation found in patient C.II.1 and the hydrocephalus-related to stenosis of the aqueduct of Sylvius and the vesico-ureteral reflux found in individual B.II.4. It remains unclear whether these isolated signs are related to *UNC45A* or not. The clinical heterogeneity could reflect differing degrees of functional deficiency resulting from distinct effects of each mutation on UNC45A expression/stability on different tissues. Although we observe similar protein expression levels in lymphoblastoid cells, we cannot exclude that the mutations could have a differential impact on specific tissues. The variable expressivity could also be impacted by modifiers, epigenetics, or environmental factors.

In conclusion, our study finds a strong association between *UNC45A* loss of function mutations and a new syndrome, Osteo-Oto-Hepato-Enteric (O2HE) syndrome. According to previous reports, *UNC45A* is possibly associated with the cytoskeleton and further evidences need to be accumulated to understand the molecular mechanisms underlying the O2HE syndrome.

**Table 1:**
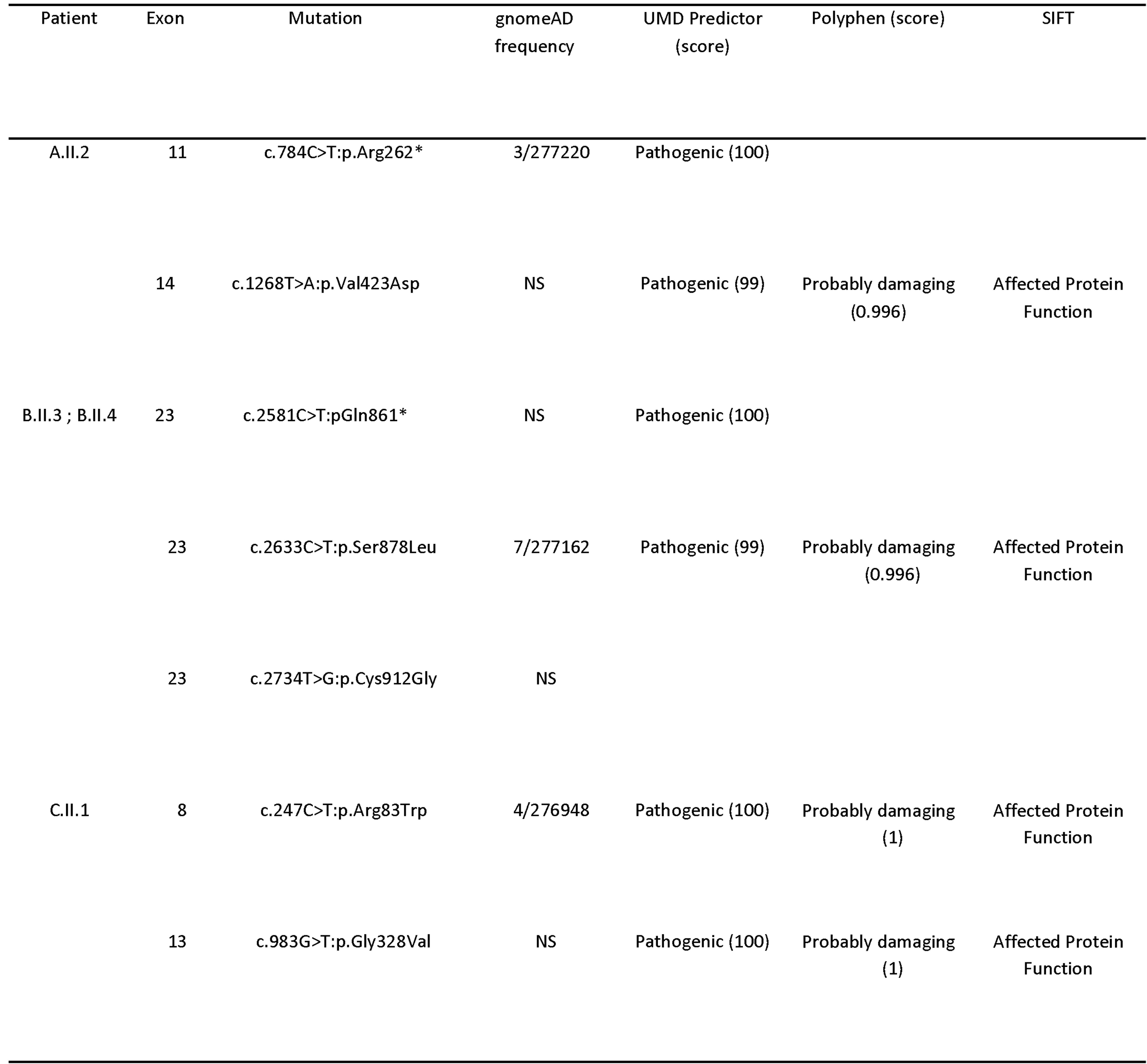
*UNC45A* mutations and pathogenicity prediction. Mutations in *UNC45A*, gnomAD (genome Aggregation Database) frequency and prediction of the pathogenicity. (NS stand for Never Seen)

## Supplemental Data

Supplemental Data include nine figures and a Supplemental Note: Case Reports.

## Acknowledgments

We are grateful to all the affected individuals and their families who participated in this study. We also thank Erica Davis and Michael Mitchell for critical reading of the manuscript and helpful suggestions and Nathalie Colavolpe for her help with X-Ray. This work was support in part by grants from the «Fondation Maladies Rares», AMGORE (« Association Méditerranéenne pour les Greffes d’ORganes aux Enfants »), the Regional Council of Burgundy (PARI 2014) and the patient support group Association Romy - La vie par un fil.

## Web Resources

UMD predictor http://umd-predictor.eu/

VarAFT http://varaft.eu.

genome Aggregation Database (c) http://www.gnomad.broadinstitute.org.

ExAC www.exac.broadinstitute.org.

1000Genomes http://www.internationalgenome.org/1000-genomes-browsers/.

PolyPhen-2 http://genetics.bwh.harvard.edu/pph2/.

Mutation Taster http://www.mutationtaster.org/.

Mutalyzer 2.0.26 https://mutalyzer.nl/.

Fidji*(Fiji Is Just ImageJ)*software

OMIM database https://www.omim.org

MAFFT mafft.cbrc.jp.

BMGE pasteur.fr/pub/GenSoft/projects/BMGE/..

NCBI https://www.ncbi.nlm.nih.gov.

CLUSTAL https://www.ebi.ac.uk/Tools/msa/clustalo/.

